# Autonomous Homeostatic Synthetic Cells via Self-Gating DNA Nanopores

**DOI:** 10.64898/2026.09.01.748542

**Authors:** Reshma Bano, Zak Marshall, Stefan Howorka, Wooli Bae

**Author notes:** Correspondence: Wooli Bae.

## Abstract

Homeostasis is a fundamental hallmark of living organisms, arising from the complex interplay between biochemical reactions and regulatory feedback systems. Reconstituting such self-regulating behaviour in minimal synthetic cells enables continuous, persistent operation of biochemical reactions for extended amount of time. In this work, we demonstrate a minimal homeostatic synthetic cell capable of autonomous flux regulation using DNA nanotechnology and bottom-up synthetic biology. Our homeostatic architecture consists of Giant Unilamellar Vesicles (GUVs) equipped with gated DNA nanopores, encapsulated *in vitro* transcription (IVT) machinery, and an RNA degradation system. We achieve homeostasis under varying external chemical stimuli specifically varying concentrations of rNTPs by implementing a negative feedback loop between rNTP influx and RNA production. In our system, DNA nanopores facilitate the influx of rNTPs from the external environment, driving internal transcription. Crucially, the transcription process generates RNA “blockers” designed to bind and gate the DNA nanopores, thereby attenuating further rNTP influx. Our system is dynamic as encapsulated RNases slowly degrade the RNA blockers, allowing the pores to reopen as blocker concentration goes down. We first characterise the functionality and gating efficiency of the DNA nanopores using both pre-synthesised and in situ produced DNA and RNA blockers. We then demonstrate that rNTP flux through these pores is sufficient to drive IVT within the GUVs. Finally, by integrating these modules, we demonstrate robust homeostasis: the system maintains a steady-state level of RNA production for up to 16 hours. By harnessing the controllability of negative feedback loop, we demonstrate thresholding of the homeostasis level using single-stranded regulator DNA. This work establishes a versatile framework for engineering adaptive and self-sustaining responsive nanomaterials and synthetic cell chassis.

## Introduction

A defining hallmark of living system is their ability to maintain internal stability in the face of challenging internal and external environment over time^1^. This dynamic property, known as homeostasis, relies on the continuous sensing and regulation of internal parameters such as pH, temperature, and molecular concentrations allowing organisms to adapt and survive^2–4^. However, in bottom-up synthetic biology, establishing chemical homeostasis for synthetic cell systems remains a significant challenge^5,6^. Most synthetic cells operate as closed systems that inevitably proceed toward thermodynamic equilibrium, severely limiting their operational lifespan and their capacity to function in continuously changing environments^7,8.^ While systems with open transport exist, their permeability is typically passive or externally controlled, falling short of the autonomous regulation characteristic of natural systems^9,10^. Designing a robust homeostatic system within a confined synthetic architecture is critical for the generation of life-like materials capable of continuous, consistent operation across diverse environments for extended period^11^.

However, achieving homeostasis in synthetic cells remains an open challenge. While buffering, and external supplementation of key ingredients can sustain biochemical reaction for extended periods, genuine homeostasis relies on negative feedback loops that synergistically integrate molecular sensing, information processing, and actuation^12–14^ for sustained biochemical reactions and modular regulation. As these machines must operate concurrently within a shared, micrometre-sized compartment, avoiding component crosstalk or irreversible kinetic trapping presents a significant engineering challenge^15^. Overcoming these limitations therefore remains an outstanding goal of the field.

Here, we demonstrate a homeostatic synthetic cell system using the programmability of nucleic acid nanotechnology^16^ and the versatility of bottom-up synthetic biology^17^. We construct giant unilamellar vesicles (GUVs)^18^ equipped with gated transmembrane DNA nanopores^19^, encapsulating in vitro transcription (IVT) machinery^20^ alongside an enzymatic RNA degradation module (Fig. 1). In our circuit, the influx of external ribonucleotide triphosphates (rNTPs) through the nanopores drives the transcription of the RNA blocker strands. The blockers act as an inhibitory feedback signal, directly binding and closing the nanopores^21^ to stop the influx of rNTPs. Simultaneously, encapsulated RNase acts as a dissipative reset, continuously degrading the RNA blockers to restore pore permeability. Through this programmed interplay of transport, transcription, gating, and enzymatic degradation, the system establishes a dynamic homeostatic equilibrium that buffers internal transcription rates at varying external rNTP concentrations over 16 hours. Furthermore, by introducing additional controller module that can supress the blockers in the form of single-stranded DNA, we demonstrate the capacity to tune the system’s steady-state threshold over a five-fold range.

**Figure 1.**
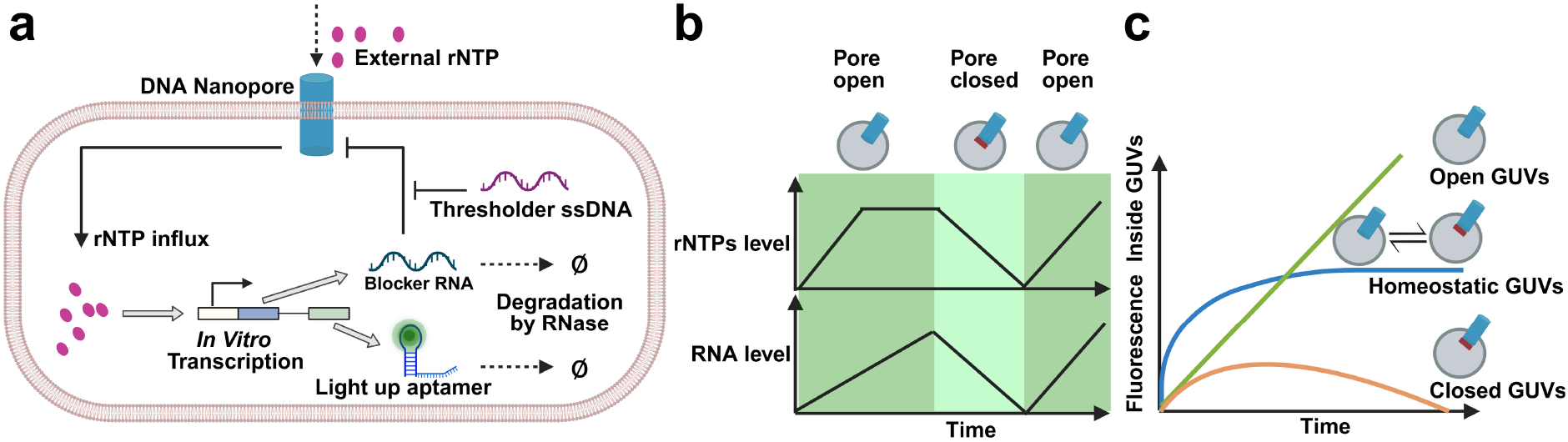
RNA-mediated homeostatic regulation in DNA nanopore-functionalized synthetic cells. **a**, Design of the homeostatic synthetic cell. DNA nanopores embedded within the lipid membrane mediate rNTP influx from the external environment. The influx of rNTPs drives *in vitro* transcription (IVT) of both blocker RNA strands that block the pores and the light-up aptamer Broccoli, which reports internal RNA levels via fluorescence. Encapsulated RNase degrades the RNA pool, allowing the pore to reopen to establish dynamic homeostasis. Single-stranded ‘thresholder’ DNA sequesters blocker RNA to modulate the RNA level at homeostasis. **b**, Time-course representation of the dynamic regulation of rNTP influx and RNA production inside a synthetic cell via DNA nanopores. In the open state, rNTPs influx drives RNA synthesis until the pore is blocked; once blocked, RNase activity depletes the internal RNA pool, prompting the pore to reopen. **c**, Expected fluorescence trajectories comparing homeostatic synthetic cells against constitutively open or closed GUVs. Permanently closed GUVs deplete their RNA, whereas permanently open GUVs accumulate RNA indefinitely.

## Results

### Characterisation of DNA nanopore-mediated molecular transport in GUVs

Sulforhodamine B (SRB) and calcein were selected as model solutes to monitor nanopore-driven efflux and influx, respectively^21^, as their molecular weights closely match those of rNTPs. GUVs were prepared via water-in-oil emulsion transfer ^18^, incorporating a cholesterol-anchored membrane DNA nanopore (Supplementary Fig. 1) based on a design by Diederichs *et al* ^19^. Confocal fluorescence imaging confirmed intact GUV formation, efficient SRB encapsulation, and the successful localisation of the nanopores to the lipid bilayer (**Fig. 2a–d**). Subsequent functional assays (**Fig. 2e,f**) revealed that both lumenal SRB leakage and external calcein influx increased by ~50% exclusively in the presence of the cholesterol-anchored DNA nanopores (**Fig. 2g,h**; Supplementary Fig. 2). The signal-to-noise ratio (SNR) was calculated from the background-corrected fluorescence intensity of individual GUVs relative to the corresponding background signal. SRB-loaded GUVs exhibited the strongest signal relative to background, whereas substantially lower SNRs were observed for without SRB and Calcein influx conditions (Supplementary Table 1).

**Figure 2.**
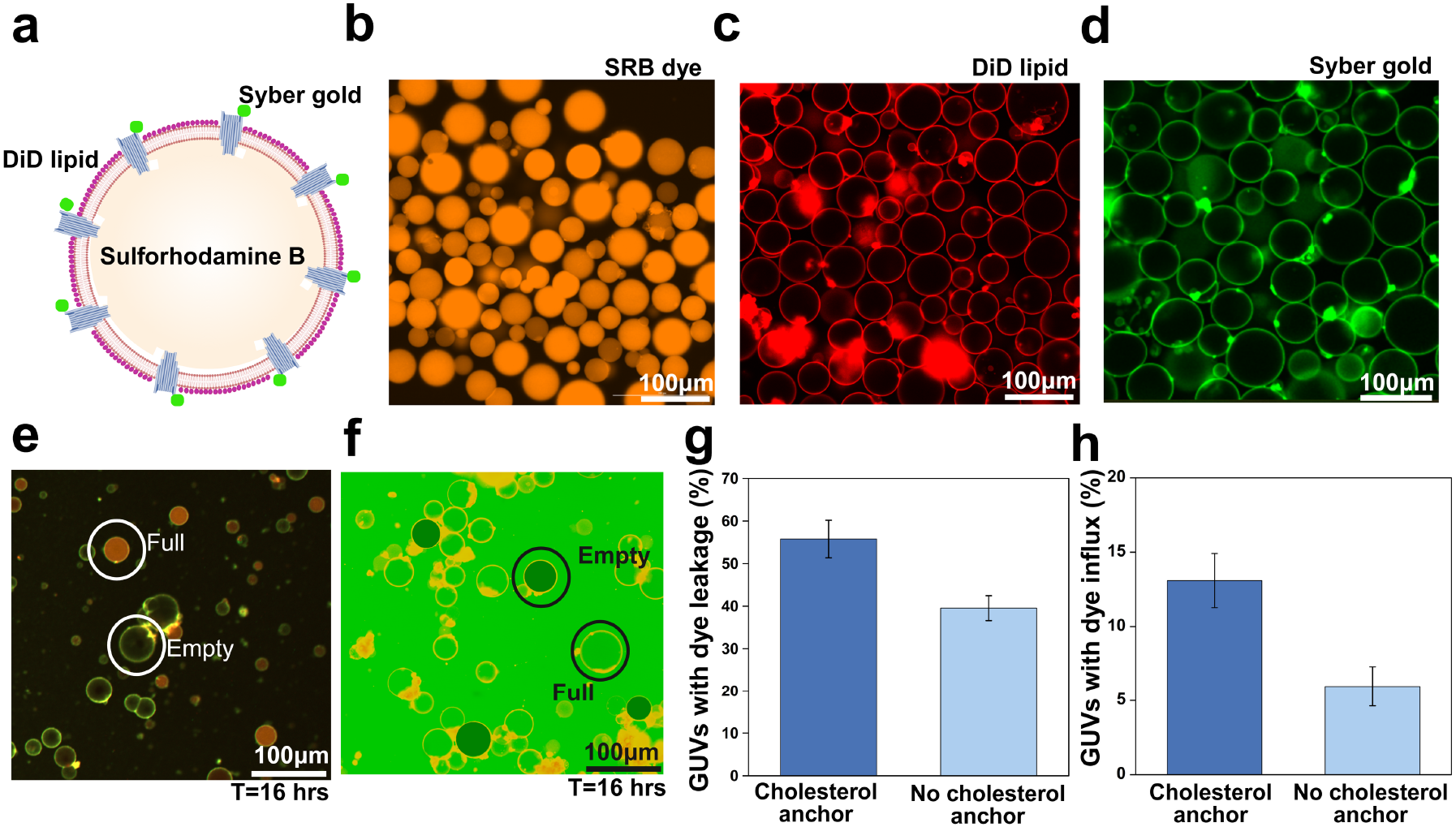
Characterisation of GUVs and DNA nanopores. **a**, Fluorescence labelling scheme for the GUV components: DiD-labelled membrane (red), encapsulated sulforhodamine B (SRB, orange), and SYBR− Gold-stained DNA nanopores (green). **b–d**, Representative confocal images of GUVs showing encapsulated SRB (**b**), DiD-labelled lipid membranes (**c**), and SYBR− Gold-stained DNA nanopores embedded within the membranes (**d**). **e, f**, Representative confocal images of nanopore-induced dye efflux (**e**) and influx (**f**) showing successful transport of small molecules mediated by the nanopores. **g, h**, Quantification of the percentage of nanopore-bearing GUVs exhibiting dye leakage and influx, measured in both the presence and absence of cholesterol-anchored DNA showing successful cholesterol-dependent activity of DNA nanopores. Data are representative of at least three independent experiments with 100-300 number of GUVs.

### Programmable control of DNA nanopore-mediated transport

Programmable control of molecular transport was achieved using DNA and RNA blocker strands (Supplementary table 2) that hybridise onto the tip of the nanopore and extends across the channel to form a molecular roadblock (**Fig. 3a**, Supplementary Fig. 3). The nanopores were blocked either prior to or following insertion into the lipid membranes by incubating with the blockers in the solution or by encapsulating the blockers inside of the GUVs. Confocal fluorescence imaging showed a marked reduction in dye influx compared with unblocked nanopores, demonstrating effective inhibition of transmembrane transport (**Fig. 3b,c**). Quantitative analysis revealed that blocking efficiency was highly comparable regardless of the blocking timing compared to the pore insertion or whether DNA or RNA blockers were used (**Fig. 3d**).

**Figure 3.**
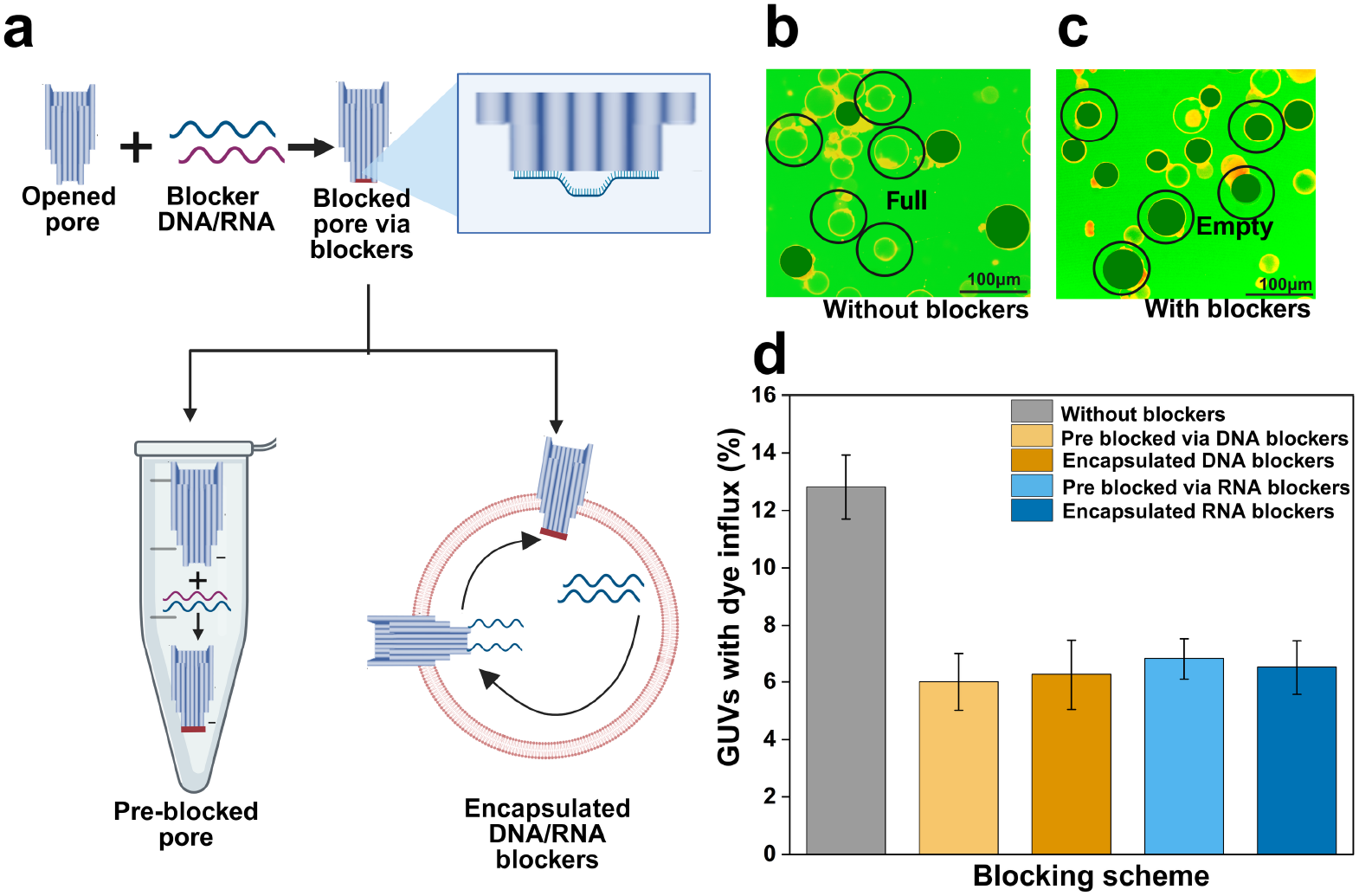
Programmable blocking of DNA nanopores using complementary oligonucleotides. **a**, Schematic illustration of nanopore gating via the DNA or RNA blocker strands. The blocking was performed either pre-insertion (in a separate reaction) or *in situ* following nanopore incorporation into the GUV membrane via encapsulated blockers. **b, c**, Representative confocal fluorescence images of dye influx in the absence (**b**) and presence (**c**) of blocker strands demonstrates the blockers can control the activity of DNA nanopores. **d**, Quantification of the percentage of GUVs exhibiting dye influx under various conditions: unblocked nanopores, pre-incubated blockers (separate reaction), and *in situ* blocking with either DNA or RNA strands showing the blockers work in every condition. Scale bars, 100 μm.

#### *In vitro* transcription inside synthetic cells driven by encapsulated or externally supplied rNTPs

As RNA production is the core metabolic activity in our system, we first verified the fluorescence response of tBroccoli RNA in the presence of DFHBI in block (Supplementary Fig. 4). We then verified whether in vitro transcription (IVT) could occur within the synthetic cell lumen using the fluorogenic RNA aptamer tBroccoli as a transcriptional reporter (Fig. 4a) also validated by using positive and negative controls (Supplementary Fig. 5). We constructed our synthetic cells by co-encapsulating the DNA template, T7 RNA polymerase, and rNTPs in GUVs. Our synthetic cells exhibited a robust increase in DFHBI fluorescence, confirming successful internal transcription (Fig. 4b).

**Figure 4.**
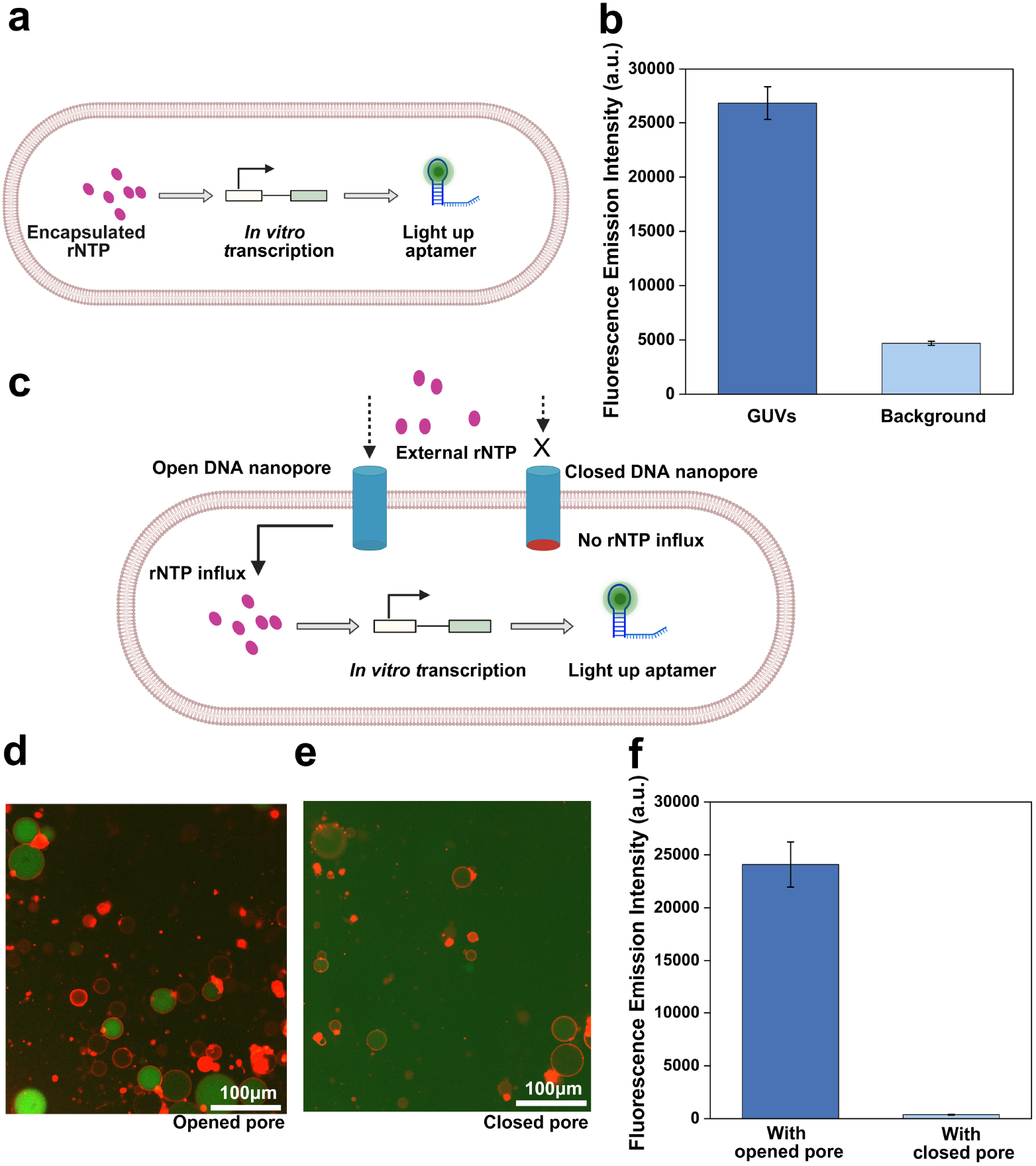
*In vitro* transcription inside synthetic cells driven by encapsulated or transported rNTPs. **a**, Schematic representation of the synthetic cells encapsulating the IVT machinery and rNTPs to transcribe light-up RNA aptamer tBroccoli for imaging. **b**, Quantification of internal DFHBI fluorescence intensity relative to the background signal, confirming successful transcription within our synthetic cells. **c**, Schematic representation of tBroccoli transcription through externally supplied rNTPs transported through open DNA nanopores; in contrast, closed nanopores prevent rNTP uptake, halting transcription. **d, e**, Representative confocal fluorescence images of synthetic cells containing open (d) and blocked (e) nanopores following incubation with externally supplied rNTPs and DFHBI. Fluorescence is observed exclusively in open-pore GUVs, whereas negligible signal is detected in blocked nanopore controls. Scale bar, 100 μm. **f**, Comparison of DFHBI fluorescence intensity inside synthetic cells bearing open versus closed nanopores showing that the rNTP transported via open nanopore can drive IVT while blocker successfully blocks influx of rNTPs and transcription.

Next, we investigated functionality of three submodules for homeostasis – sustained IVT via external rNTPs, gating of the nanopores via internally produced RNA and dissipating of the system via RNase. First, we constructed synthetic cells encapsulating IVT machineries except for rNTPs. Following incubation with 8 mM of external rNTPs, synthetic cells equipped with open DNA nanopores generated a strong tBroccoli fluorescence signal, whereas negligible fluorescence was observed in synthetic cells with blocked nanopores (Fig. 4c–f). Then we performed permeability test for DNA nanopores for synthetic cells transcribing RNA blockers internally via encapsulated IVT machineries. This demonstrated, successful autonomous gating of DNA nanopores on synthetic cells for small-molecule transport (Supplementary Fig. 6). Finally, we performed RNase screening for controlled degradation and turnover of RNA blockers and reporters (Supplementary Fig. 7).

### Demonstration of homeostasis and thresholding

To achieve homeostatic synthetic cells, we combined the individual submodules by encapsulating IVT machinery, DNA templates for RNA blockers, and an RNase mix inside GUVs. We immobilised the synthetic cells on a 96-well plate and introduced DNA nanopores alongside external rNTPs ranging from 4–8 mM (Figure 5a). Time-lapse confocal microscopy confirmed continuous tBroccoli RNA expression. As anticipated, the tBroccoli signal reached a steady state, despite having both production and degradation machineries, confirming intracellular RNA homeostasis (Figure 5b) (Supplementary Fig. 8). This signal remained stable for 16 hours, demonstrating robust homeostasis despite continuous RNA turnover via RNases (Figure 5c,d). In contrast, synthetic cells containing opened DNA nanopores or closed without pore (Figure 1c) failed to establish RNA homeostasis. Permanently opened nanopores resulted in continuous RNA accumulation, whereas permanently closed membrane prevented sustained RNA production, demonstrating that homeostasis emerges only through the dynamic feedback between nanopore opening and RNA-mediated pore closure (Figure 5e).

**Figure 5.**
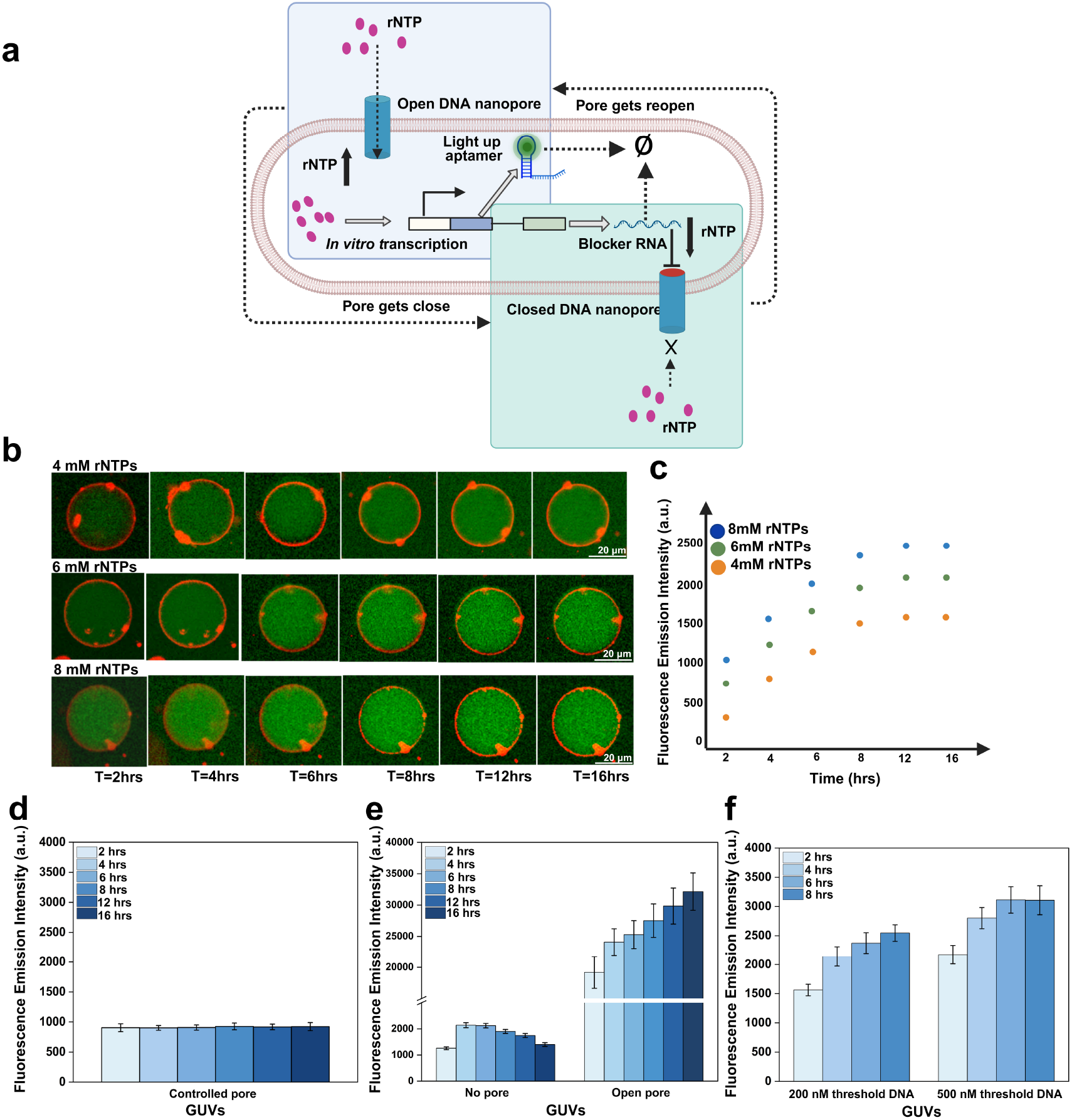
Homeostatic regulation and threshold inside synthetic cells by reversible DNA nanopore gating system. **a**, Schematic representation of the homeostatic circuit. IVT-generated blocker RNA closes the DNA nanopores, whereas RNase-mediated blocker degradation reopens them, enabling repeated control of rNTP influx. **b**, Representative confocal images of synthetic cells supplied with 4, 6 or 8 mM rNTPs at different time point up to 16 hours. Scale bar, 20 μm. **c**, Exemplar time-dependent fluorescence intensity inside synthetic cells corresponding to confocal images supplied with external rNTPs, showing concentration-dependent signal increase and stabilisation over time. **d**, Average fluorescence intensity of homeostatic synthetic cells with time points showing homeostasis up to 16 hours. **e**, Synthetic cells with closed membrane or constitutively opened DNA nanopores as illustrated in Figure 1c. Fluorescence signal from closed synthetic cells show initial increase followed by continuous decline due to degradation by RNase while open synthetic cells exhibited a continuous increase fluorescence signal over 16 h. **f**, Modulation of RNA expression by 200 nM or 500 nM threshold antisense DNA in homeostatic synthetic cells showing the higher RNA expression.

To further highlight the versatility of this feedback network, we introduced a simple single-stranded antisense DNA designed to bind and deactivate the blocker RNA. This inhibitor acts as a threshold by sequestering a defined amount of RNA prior to nanopore blockage. As a result, the synthetic cells successfully achieved RNA homeostasis at tuned, elevated levels corresponding to the concentration of the inhibitor DNA at 200 nM and 500 nM (Figure 5f).

## Discussion

Here we establish nanopore-mediated negative feedback as a general strategy for engineering autonomous homeostasis in synthetic cells. Although membrane transport, compartmentalised gene expression and programmable DNA nanostructures have each been demonstrated previously, these components have largely functioned independently, limiting their ability to dynamically regulate intracellular biochemical states^22–25^. By coupling programmable DNA nanopores with in vitro transcription and RNA degradation, we integrate membrane transport directly into a gene-expression feedback circuit that autonomously maintains stable intracellular RNA levels. Rather than serving solely as a physical boundary, the membrane therefore becomes an active regulatory interface governing intracellular molecular homeostasis inferring information from intracellular state^26^.

An important advantage of our system is the programmability. DNA nanopores are transformed from passive transport channels into regulatory elements whose permeability is determined by intracellular molecular information^27^. The homeostatic operating point can be quantitatively tuned through thresholder DNA without redesigning the nanopore itself, providing a simple mechanism for programming intracellular RNA concentrations. Because blocker sequences are independently programmable^28–30^, membrane transport could also be coupled to functional RNAs, including aptamers, riboswitches and catalytic RNAs^31–34^, allowing transport to respond selectively to specific biochemical signals including proteins.

The prolonged operation of the homeostatic circuit also illustrates the advantages of biochemical compartmentalisation^35,36^. Encapsulation of T7 RNA polymerase and RNase within lipid vesicles is expected to preserve enzymatic activity by protecting the enzymes from surface adsorption and extracellular proteolytic degradation, as demonstrated previously for compartmentalised cell-free systems^37–39^. In parallel, feedback-controlled nanopore gating restricts substrate influx once the homeostatic state has been established, providing a potential mechanism for improving substrate utilisation during long-term cell-free reactions. Together, these features enable sustained autonomous regulation while maintaining continuous communication between the synthetic cell and its external chemical environment.

The design principles demonstrated here should be readily extended beyond RNA homeostasis. Coupling membrane transport to functional RNAs with defined regulatory activities would enable transport to respond directly to intracellular biochemical signals, while the same feedback architecture could be expanded to protein synthesis by linking nanopore permeability to protein-mediated regulation of transcription or transport. More broadly, integrating programmable membrane transport with intracellular reaction networks provides a modular strategy for constructing synthetic cells capable of maintaining stable internal states while adapting their biochemical activities to changing environmental conditions. Such transport-mediated regulatory circuits represent an important step towards increasingly autonomous, adaptive and life-like synthetic cells.

## Materials and Methods

### 1. DNA nanopore folding, purification and characterisation

All DNA staple strands, blocker DNA strands, sequester DNA strands and 3' cholesterol-modified DNA oligonucleotides were purchased from Integrated DNA Technologies (IDT). The 5' cholesterol-modified DNA strand was purchased from Eurogentec. Single-stranded M13mp18 scaffold DNA was obtained from Tilibit Nanosystems. DNA nanopores were assembled from a previously reported caDNAno design^19^. DNA nanopores were folded by mixing 20 nM M13mp18 scaffold DNA with approximately 200 nM of each staple strand in folding buffer containing 1× TAE (cat#1061741000, Merck millipore) and 25 mM MgCl_2_ (cat# J61014, Thermo Fisher Scientific). The folding mixture was subjected to thermal annealing consisting of an initial incubation at 85°C for 20 s, followed by cooling from 65°C to 40°C at a rate of −0.8 °C h^−1^, further cooling from 40°C to 4°C at −2.0 °C h^−1^, and storage at 4 °C after completion of the annealing programme. Folded DNA nanopores were purified using polyethylene glycol (PEG) precipitation method^40^. Equal volumes of the folding reaction and precipitation buffer (15% (w/v) PEG 8000 (cat#043443.A3, Thermo Fisher Scientific), 10 mM Tris (cat#AM9855G, Thermo Fisher Scientific), 1 mM EDTA (cat#AM9855G, Thermo Fisher Scientific)) and 505 mM NaCl (cat#AM9855G, Thermo Fisher Scientific) were mixed thoroughly by gentle inversion and centrifuged at 16,000 × g for 25 min at 4 °C. Following centrifugation, the supernatant was carefully removed, and the DNA nanopore pellet was resuspended in outer aqueous solution (OAS) consisted of 300 mM glucose (Cat#15023021, Thermo Fisher Scientific), 20 mM HEPES (cat#15630056, Thermo Fisher Scientific), 10 mM MgCl_2_ and 100 mM KCl (cat# J63739.AK, Thermo Fisher Scientific) before using storage at −20 °C. The assembled DNA nanopores was assessed by agarose gel electrophoresis. Samples were analysed on 2.0% agarose gels (cat# A9539, Sigma-Aldrich) containing 1× SYBR Safe stain (cat# S33102, Thermo Fisher Scientific) in 1× TAE buffer supplemented with 10 mM MgCl_2_. Electrophoresis was performed at 90 V for 120 min at 4 °C, after which the gels were visualised under blue-light illumination.

### 2. GUVs preparation

Giant unilamellar vesicles (GUVs) were prepared using the inverted emulsion phase-transfer method. Lipid-in-oil solution was prepared from 1,2-dioleoyl-sn-glycero-3-phosphocholine (18:1 DOPC; Cat#850375) and 1,2-dioleoyl-sn-glycero-3-phosphoethanolamine-N-(biotinyl) (sodium salt) (18:1 Biotinyl DOPE; Cat#870282) (Avanti Polar Lipids), together with the lipid dye 1,1'-dioctadecyl-3,3,3',3'-tetramethylindodicarbocyanine, 4-chlorobenzenesulfonate salt (DiD; Cat#D7757, Thermo Fisher Scientific). Stock solutions prepared in chloroform or ethanol were stored at −20 °C before use. Lipids were mixed in 20 mL WHEATON liquid scintillation vials (Cat#DWK986546, Sigma-Aldrich) at a weight ratio of 99.8:0.1:0.1 (18:1 DOPC:18:1 Biotinyl DOPE) using stock concentrations of 25 mg mL^−1^, 1 mg mL^−1^ and 2 mg mL^−1^, respectively. Organic solvents were removed under a gentle stream of nitrogen while continuously rotating the vial to form a thin lipid film. The dried lipid film was resuspended in 2 mL BioReagent mineral oil (Cat#M5904, Sigma-Aldrich) to obtain a final lipid concentration of 2 mg mL^−1^. The vials were sealed with Parafilm and sonicated in a 40°C water bath for 30 min. Lipid-in-oil solutions were freshly prepared and used on the day of the experiment. Water-in-oil emulsions were generated by mixing the internal aqueous solution (IAS) at 2.5% (v/v) with 400 μL of the lipid-in-oil solution using the rumble-strip method (approximately one pass per second for ten passes). Subsequently, 100 μL of the emulsion was carefully layered onto the outer aqueous solution (OAS) contained in a 1.5 mL microcentrifuge tube and centrifuged at 3,000 × g for 10 min at room temperature to transfer the emulsion droplets across the oil–water interface, thereby forming GUVs. The resulting GUV pellet was collected by gentle aspiration and transferred into a fresh microcentrifuge tube for subsequent experiments. For dye leakage and influx assays, the IAS consisted of 300 mM sucrose (Cat#A15583.36, Thermo Fisher Scientific), 20 mM HEPES, 10 mM MgCl_2_ and 100 mM KCl prepared in nuclease-free water. Sulforhodamine B (SRB; Cat#S1402, Sigma-Aldrich) was included at a final concentration of 50 μM for dye leakage experiments only. The OAS consisted of 300 mM glucose (Cat#15023021, Thermo Fisher Scientific), 20 mM HEPES, 10 mM MgCl_2_ and 100 mM KCl. Calcein was added to the OAS at a final concentration of 50 μM for dye influx experiments only. A volume of 100 μL OAS was used during GUV preparation.

### 3. In vitro transcription in bulk and inside GUVs

In vitro transcription (IVT) was first performed in bulk to confirm DFHBI fluorescence upon binding to the transcribed tBroccoli RNA aptamer. Following validation in bulk, IVT was performed inside GUVs to assess DFHBI fluorescence resulting from tBroccoli transcription within the vesicles. For IVT experiments in GUVs, the inner aqueous solution (IAS) consisted of 300 mM sucrose, 100 mM KCl, 20 mM HEPES, 10 mM MgCl_2_, IVT buffer, T7 RNA polymerase, rNTPs, DNA templates encoding tBroccoli or RNA blocker strands, RNase inhibitor and 50 μM DFHBI prepared in nuclease-free water. The corresponding outer aqueous solution (OAS) contained 300 mM glucose, 100 mM KCl, 20 mM HEPES, 10 mM MgCl_2_, IVT buffer and 50 μM DFHBI. A 50-μl volume of OAS was used during GUV preparation. For homeostasis experiments, RNase titration was first performed in bulk to determine the appropriate RNase concentration and assess its effect on RNA degradation. Based on these bulk measurements, RNase inhibitor in the IAS was replaced with RNase, whereas rNTPs were omitted from the IAS and supplied exclusively through the OAS.

### 4. Preparation of passivated surfaces for GUV immobilisation

To minimise GUV adhesion and membrane rupture during imaging, 96-well microplates (Cat#165305, Thermo Fisher Scientific) were passivated before use. Each well was incubated with 200 μL of 5 mg mL^−1^ bovine serum albumin (BSA; Cat#A9647, Sigma-Aldrich) prepared in 1× PBS and supplemented with 0.5% biotin-labelled bovine serum albumin (Bio-BSA; Cat#A8549, Sigma-Aldrich) for 30 min at room temperature. Following incubation, the wells were washed four times with 1× PBS (Cat#AM9625, Thermo Fisher Scientific). After the final washing step, 200 μL of 20 ng μL^−1^ NeutrAvidin (Cat#31055, Thermo Fisher Scientific) was added to each well and incubated for 30 min at room temperature. The NeutrAvidin solution was then removed, leaving approximately 50 μL in each well, and the wells were subsequently washed six times with 150 μL of outer aqueous solution (OAS). Passivated microplates were used on the day of preparation.

### 5. Confocal Microscopy

Following preparation, GUVs were recovered by aspirating the oil phase and resuspending the GUV pellet (A total of 30 μL of the solution containing GUVs) in passivated and immobilisation-treated 96-well microplates. Cholesterol-modified DNA anchor strands were added to the GUV suspension at a final concentration of 200 nM and incubated for 10 min to allow membrane insertion. Purified folded DNA nanopores were subsequently added to a final concentration of 3 nM and incubated for an additional 15 min before imaging. Confocal fluorescence imaging was performed using a BioTek Cytation C10 Confocal Imaging Reader (Agilent Technologies) equipped with a 20× objective and a 60 μm spinning-disk confocal module. Fluorescence signals from the membrane dye, encapsulated fluorophores and membrane-incorporated DNA nanopores were acquired using the GFP, RFP and Cy5 channels, respectively. To enable quantitative comparison of fluorescence intensities across experiments, all images were acquired using a fixed detector gain, and pixel intensities were normalised to a standard exposure time of 25 ms based on the integration time used for each acquisition. Images were exported in both TIFF and PNG formats for subsequent processing and analysis. Each experimental condition was independently repeated between three and five times.

### 6. Image processing and quantitative analysis

Confocal fluorescence images were analysed using Fiji: an open-source platform for biological-image analysis. All images within each experiment were processed using identical display and analysis settings. Fluorescence intensities were quantified by manually defining regions of interest (ROIs) around individual GUVs and measuring the mean fluorescence intensity. Background fluorescence was measured from adjacent regions lacking GUVs and subtracted from the corresponding vesicle intensity. The signal-to-noise ratio (SNR) was calculated from the background-corrected fluorescence intensity of each GUV relative to the corresponding background signal. For dye leakage and dye influx assays, GUVs were identified following contrast enhancement in fiji (ImageJ) to facilitate vesicle detection. Vesicles were visually counted and classified according to the presence or absence of detectable fluorescence leakage or influx using identical analysis criteria across the experimental conditions. The percentage of GUVs exhibiting dye leakage or influx was calculated by dividing the number of positive vesicles by the total number of GUVs analysed for each sample.

