## Supplementary material for "Autonomous Homeostatic Synthetic Cells via Self-Gating DNA Nanopores": SI figures and tables

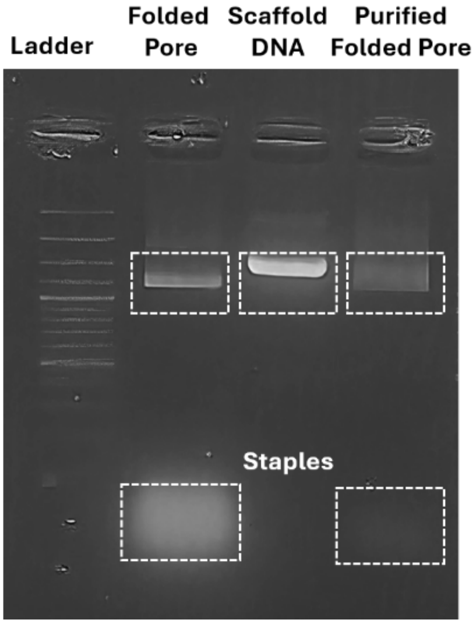


**Figure S1. Confirmation gel of DNA nanopore after folding and PEG purification.**

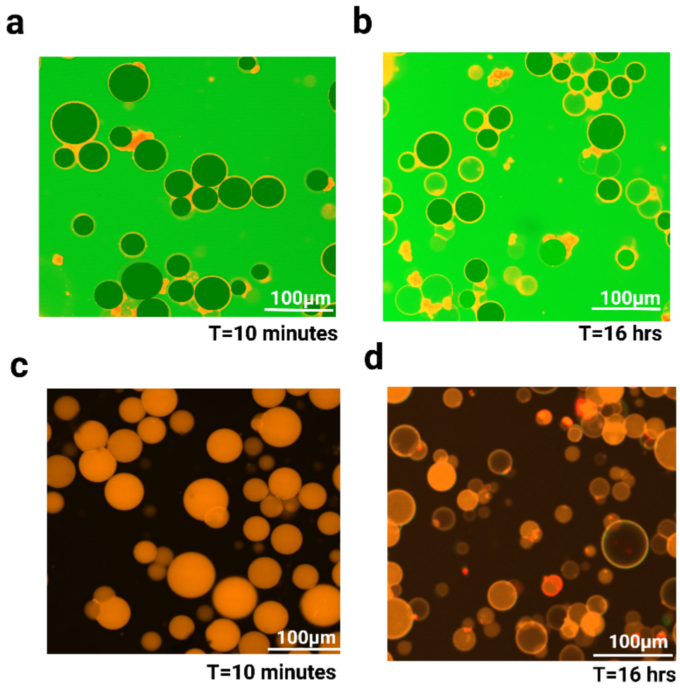


**Figure S2. Confocal imaging of GUVs incubated with DNA nanopores lacking cholesterol anchors.**

Representative confocal images showing dye leakage and influx after 16 hrs incubation of GUVs with DNA nanopores without cholesterol anchors.

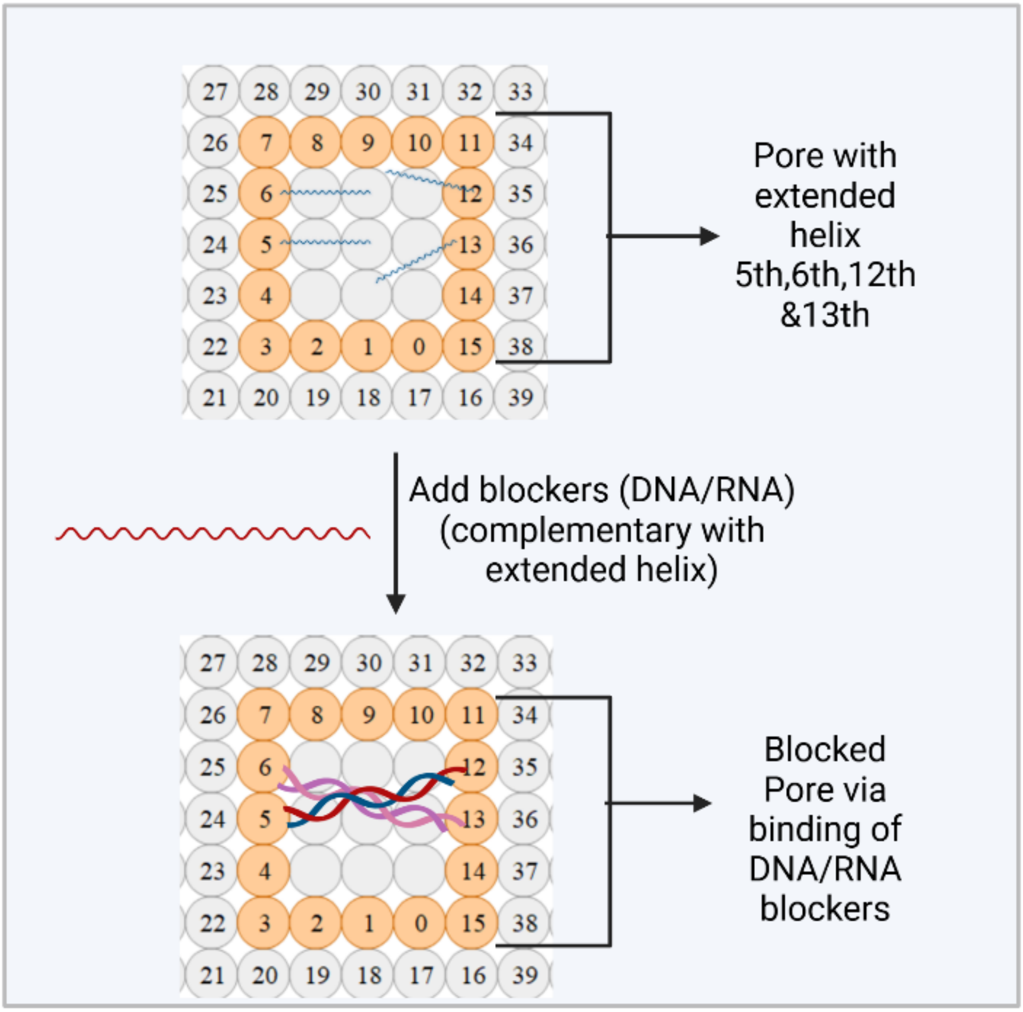


**Figure S3. Blocking scheme for DNA nanopore gating.**

Four helices of the DNA nanopore were extended to provide binding sites for complementary blocker strands. The blocker strands hybridize with the extended helices to sterically occlude the nanopore and restrict molecular transport.

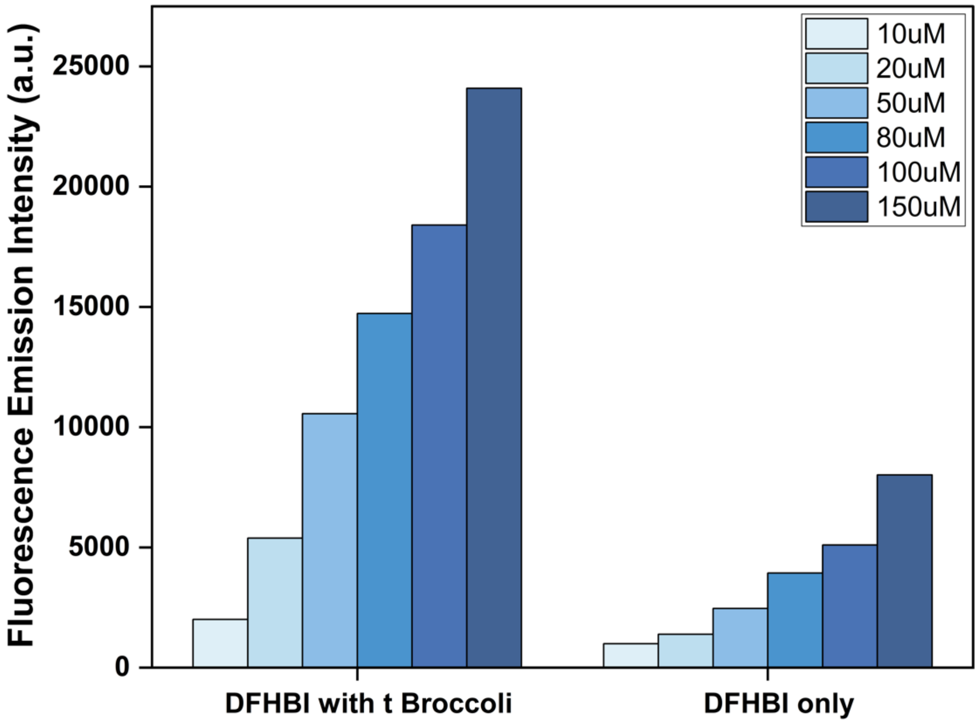


**Figure S4. tBroccoli RNA aptamer–DFHBI fluorescence in blocks.**

Quantitative analysis of DFHBI fluorescence in blocks in the presence or absence of tBroccoli RNA.

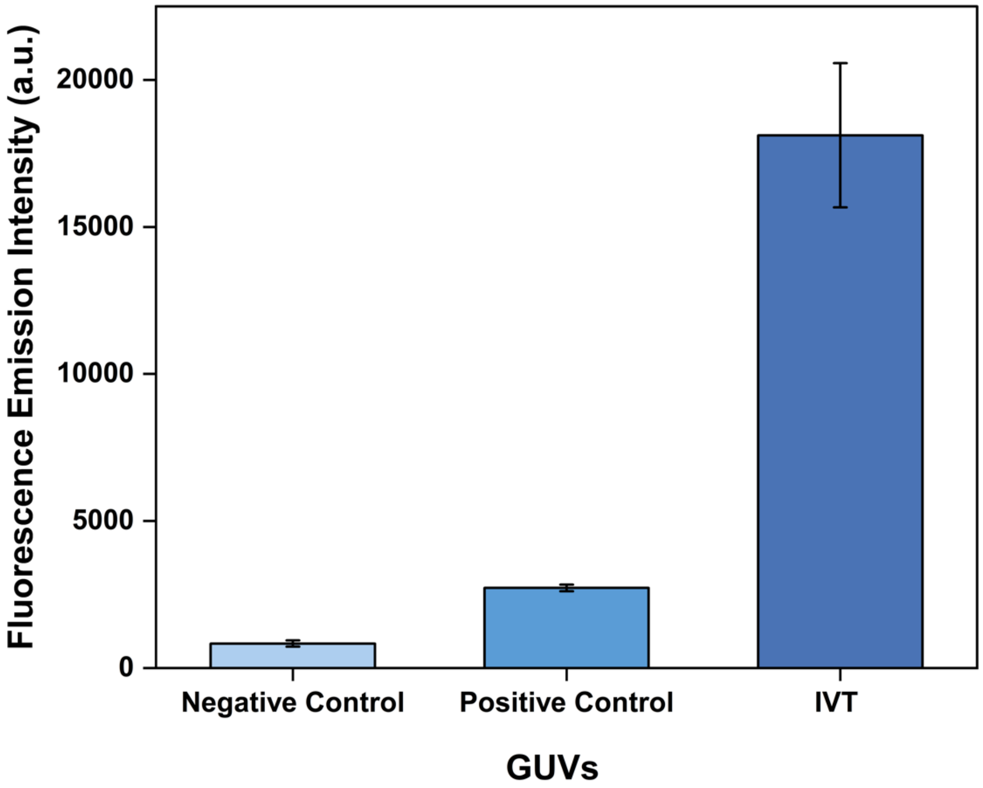


**Figure S5. IVT of tBroccoli RNA within GUVs.**

Quantitative analysis of DFHBI fluorescence in GUVs under different experimental conditions: negative control (without tBroccoli RNA) positive control (encapsulated tBroccoli RNA with DFHBI), and in situ IVT of tBroccoli using encapsulated transcription machinery, tBroccoli template DNA and DFHBI. Fluorescence from the IVT condition indicates successful transcription of tBroccoli RNA within the GUV lumen.

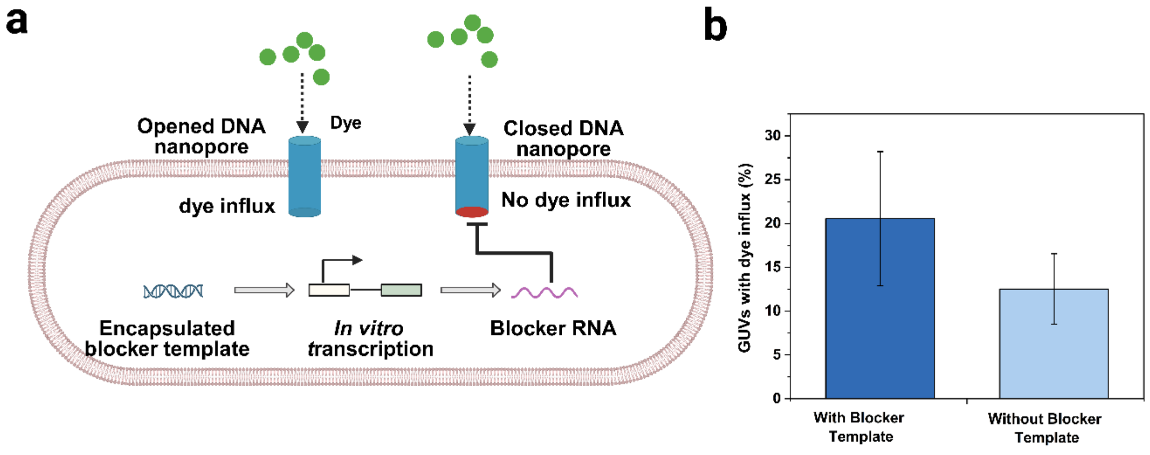


**Figure S6. Autonomous gating of DNA nanopores in GUVs.**

a, Schematic representation of autonomous DNA nanopore gating in synthetic cells for small-molecule transport using a blocker-producing template. b, Quantitative analysis of dye influx into GUVs containing IVT reactions with or without the blocker-producing template.

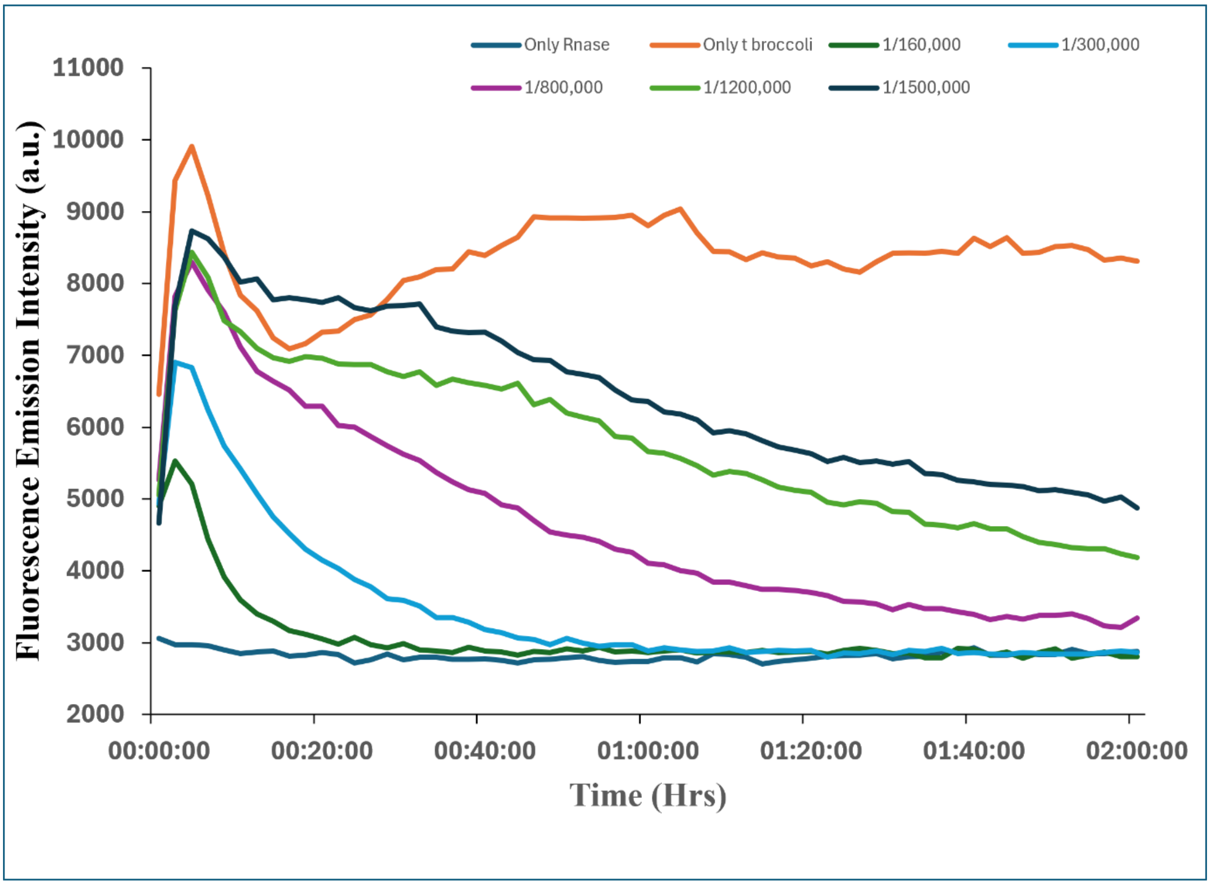


**Figure S7. RNase titration in block experiment.**

Concentration-dependent degradation of tBroccoli RNA by RNase within the blocks, assessed by the corresponding decrease in DFHBI fluorescence intensity.

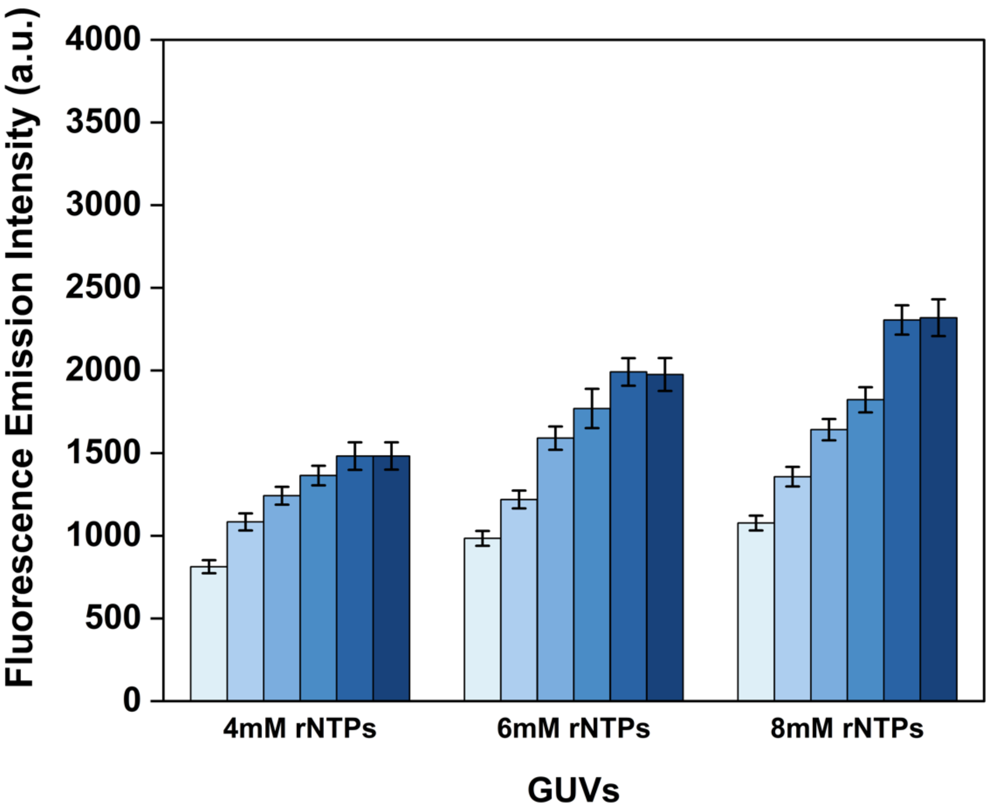


**Figure S8. Homeostatic regulation inside synthetic cells by reversible DNA nanopore.** Average fluorescence intensity of homeostatic synthetic cells at the various rNTPs concentrations and time points showing homeostasis up to 16 hours.

**Table S1. SNR table**

| GUVs | Signal to noise ratio (SNR) |
|---|---|
| With SRB dye (Full) | 44.4 |
| Outflux of SRB dye (Empty) | 5.7 |
| Without Calcein dye (Empty) | -13.1869 |
| Influx of Calcein dye (Full) | 0.18 |

**Table S2. DNA Blockers, Blocker's producers, Sequester's sequences, tBroccoli's sequences, tBroccoli with blocker producer**

| Name | Sequences 5' to 3' |
|---|---|
| Blocker 1 | ATACGTCTTCGTCGGTACCCGGACCTTGCGTCGATCATCCACGACAGAACAAGTGT<br>CGCTTCGAGCTCTACCTCCTTTGATC |
| Blocker 2 | ATACATCTGAGGGTACCCAAGCAGGGAGTCAGAACAGGGCCCAGCCCGGTCGGG<br>TACACTGTTCGGTTCACTGACAGGCAAG |
| Blocker 1 producer template | AAAAATGGCAAGGTACTCACTAATACGACTCACTATAGATACGTCTTCGTCGGTACCC<br>GGACCTTGCGTCGATCATCCACGACAGAACAAGTGTCGCTTCGAGCTCTACCTCCTT<br>TGATCAGTACTGCTAGTACGCTATTCTTTAGCAAAAA |
| Blocker 2 producer template | AAAAATGGCAAGGTACTCACTAATACGACTCACTATAGATACATCTGAGG<br>GTACCCAAGCAGGGAGTCAGAACAGGGCCCAGCCCGGTCGGGTACACT<br>GTTCGGTTCACTGACAGGCAAGAGTACTGCTAGTACGCTATTCT<br>TTAGCAAAAA |
| Sequester 1 | GATCAAAGGAGGTAGAGCTCGAAGCGACACTTGTTCTGTCGTGGATG<br>ATCGACGCAAGGTCCGGGTACCGACGAAGACGTAT |

| | |
|---|---|
| Sequester 2 | CTTGCCTGTCAGTGAACCGAACAGTGTACCCGACCGGGCTGGGCCCTGTTCTGAC<br>TCCCTGCTTGGGTACCCTCAGATGTAT |
| Extended helix 5th | GGTAGAGCTCGAAGCGACACTGTCGCTGAGCCCACGCATAA |
| Extended helix 6th | GAGGCTTTGGAACGAGGGTATGACTCCCTGCTTGGGTACCC |
| Extended helix 12th | GAATACACTAGATAAATTGTGTCGAACTCCATGTACTTTGAAACGCAAGGTCCGGGT<br>ACCGAC |
| Extended helix 13th | AGTGAACCGAACAGTGTACCCAACGGTGTTCAACGTAACAA |
| tBroccoli | CGCCGCTAATACGACTCACTATAGGCCCGGATAGCTCAGTCGGTAGAGCAGCGG<br>AGACGGTCGGGTCCAGATATTCGTATCTGTCGAGTAGAGTGTGGGCTCCGCGGGTC<br>CAGGGTTCAAGTCCCTGTTCGGGCGCCAGGCGGGCTTTTCTGGTACACG |
| tBroccoli with blocker1 | CGCCGCTAATACGACTCACTATAGGATACGTCTTCGTCGGTACCCGGACCTTGCGTC<br>GATCATCCACGACAGAACAAGTGTCGCTTCGAGCTCTACCTCCTTTGATCTTTTTGC<br>CCGGATAGCTCAGTCGGTAGAGCAGCGGAGACGGTCGGGTCCAGATATTCGTATCT<br>GTCGAGTAGAGTGTGGGCTCCGCGGGTCCAGGGTTCAAGTCCCTGTTCGGGCGCC<br>AGGCGGGCTTTTCTGGTACACG |
